# First report of a successful development of yam hybrids (*Dioscorea alata* L.) from lyophilized and long-term stored pollens

**DOI:** 10.1101/2023.03.12.532272

**Authors:** Erick Malédon, Elie Nudol, Christophe Perrot, Marie-claire Gravillon, Ronan Rivallan, Denis Cornet, Hâna Chair, Komivi Dossa

**Affiliations:** CIRAD, UMR AGAP Institut, 97170 Petit Bourg, Guadeloupe, France; UMR AGAP Institut, Univ Montpellier, CIRAD, INRAE, Institut Agro, F-34398 Montpellier, France; CIRAD, UMR AGAP Institut, F-34398 Montpellier, France

**Keywords:** Yam, freeze-drying, pollen conservation, genetic gain

## Abstract

**Background:** Greater yam, *Dioscorea alata* L., is a significant food security crop in tropical areas. However, low genetic diversity and various biological constraints, including susceptibility to viruses, ploidy, erratic and low flowering intensity, and asynchronous flowering hinder successful hybrid development and genetic gains in greater yam breeding programs. Therefore, pollen storage has gained much attention to facilitate genetic material exchanges, artificial pollinations and to increase the genetic gains in breeding programs. This 4-year study aimed at developing a practical long-term pollen storage technique for the successful development of yam hybrids. Fresh pollens were collected from two *D. alata* males, then lyophilized (two lyophilization treatments were applied), followed by storage at room temperature (24-25 °C) for 12 months. Moreover, the lyophilized and stored pollens were tested for viability by crossing with four female varieties.

**Results:** Our results showed that lyophilization is effective for achieving viable pollens after 12 months of storage. Treatment 1 (48 h drying) showed higher pollen germination and fertility rates than Treatment 2 (72 h drying). Although we observed a reduction in viability of lyophilized pollens after 12 months of storage, we generated hybrid seedlings with success rates from 12 to 21% compared to 21-31% when using fresh pollens. Paternity testing based on molecular genotyping confirmed the hybrid status of the obtained seedlings, which grew well in a greenhouse.

**Conclusions:** The results signify the importance of pollen lyophilization for yam breeding programs.

## Background

Yams are considered food security crops and are widely consumed for nutritionally rich tubers, particularly in developing countries [1–4]. Aside from the nutritional composition, including carbohydrates, proteins, minerals, and vitamins, yams play a key role in social, cultural, and religious aspects of lifestyle in western Africa [5]. Moreover, yams are rich in secondary metabolites such as saponins, diterpenoids, and alkaloids [6–8]. Over the years, progress has been made in exploiting the available genetic resources for improving the nutritional quality, and adaptability of yam species. However, the complex genetic architecture of yam species as well as numerous biotic and abiotic factors hinder further breakthroughs, offering new avenues for research and development [9].

*Dioscorea alata* L., also known as greater yam, is among the important cultivated yam species from the *Dioscorea* genus. It is a diecious species widespread across tropical areas of Africa, Asia, Oceania, and South America [10]. Breeding programs significantly contribute to crop improvement, followed by commercial propagation. However, very limited progress has been achieved so far in greater yam breeding programs [9]. The genetic diversity in *D. alata* germplasms available to breeders is very low [10, 11]. Hence, exchange of genetic materials among breeding programs is essential to make substantial genetic gains. However, exchanging yam genetic materials is very complicated because yams are naturally prone to diverse viruses and the sanitation process to achieve virus free vitroplants is a complex and time-consuming task [12, 13]. Besides, several biological constraints (polyploidy, erratic flowering, asynchronous flowering, inter-and intra-genotypic incompatibility, long vegetative growth period, low multiplication ratio, and high heterozygosity) impede varietal improvement and genetic gains in *D. alata* [10,14–19]. These constraints limit the number of compatible fertile parents, resulting in a low number of successful crosses. *Dioscorea alata* is an erratic flowering dioecious plant. Mostly the female flowers mature well before the male flowers, thus making pollination difficult between two plants [19–22]. Moreover, yam pollen is fertile for a very short time (less than 24h) at ambient temperature, complicating the crossing procedure. Therefore, pollen storage has gotten much attention to attain the maximum number of crosses during the short flowering period of female greater yam varieties. Stored yam pollens can be exchanged safely among breeding programs to increase the gene pool and accelerate the genetic gains.

Free-preserved pollen is often used to achieve synchronized flowering for a successful crossing program [23–26]. Cryopreservation allows pollen to remain viable over a long period but requires sophisticated equipment [27–29]. The pollen viability under cryopreservation is highly dependent on a stable temperature [28]. Although some studies have demonstrated the viability of cryopreserved pollen in yams [21, 24, 27], there is an apparent lack of evidence for the successful development of hybrids from long-term cryopreserved pollen. Lyophilization (freeze-drying) is an alternative method for long-term storage, consisting of freezing, sublimation, and adsorption [30]. The pollen, thus dried, can be stored in an ambient atmosphere without keeping it frozen. Previous reports suggested a successful pollen lyophilization for long-term pollen storage in different crops, for instance, sorghum [31], eucalyptus [32], maize [33,34], and pigeonpea [35]. However, lyophilization has never been tested in greater yam for a successful hybrid development. Moreover, considering that yams are mainly produced in the developing world, most laboratories in these regions do not have the facilities to stably preserve the pollen at −80 °C for a long period. It is, therefore, necessary to find alternative and practicable pollen conservation methods for yam breeding programs in developing countries.

This study aimed at testing the lyophilization approach for long-term pollen storage in *D. alata.* Moreover, we seek to evaluate the ability of the long-term stored pollens to successfully develop hybrid yam plants.

## Methods

### Plant materials, flower sampling and pollen collection

This study was conducted over four years in Guadeloupe at the Centre de coopération Internationale en Recherche Agronomique pour le Développement (CIRAD), research station of Roujol. Two *D. alata* male varieties (14M, and Toufi-Tetea) and four female varieties (74F, Boutou, Ti-violet and CT198) were obtained from the germplasm collection of CIRAD, Guadeloupe. Varieties 14M, Boutou and 74F are diploid, while Toufi-Tetea, Ti-violet and CT198 are tetraploid [19]. The male varieties were planted over three years (2019-2021) in order to collect and store pollen for the pollination of female varieties in consecutive years. The female varieties were planted in 2020 (74F, and CT198) and 2021 (74F, Boutou, Ti-violet and CT198) at the CIRAD research station of Roujol, in Guadeloupe (16°10’ 56” N, 61° 35’ 24’’ W, 10 meters above sea level). The flowering of *D. alata* begins with female varieties in Guadeloupe. The first inflorescences are visible around mid-September. Usually, around mid-October, male inflorescences are observed. Full flowering begins towards the end of October. We have noticed that the male tetraploid inflorescences are very close to anthesis from the first hours of the morning around 9 am, while the male diploid flowers do not open until around 11 am. The male flowers were collected from the male varieties by the end October each year when the petals are opened (Figure 1), corresponding to the period of maximum pollen fertility [36]. In the laboratory, the fully opened spike flowers were taken from the inflorescences and stored in Eppendorf tubes.

**Figure 1.**
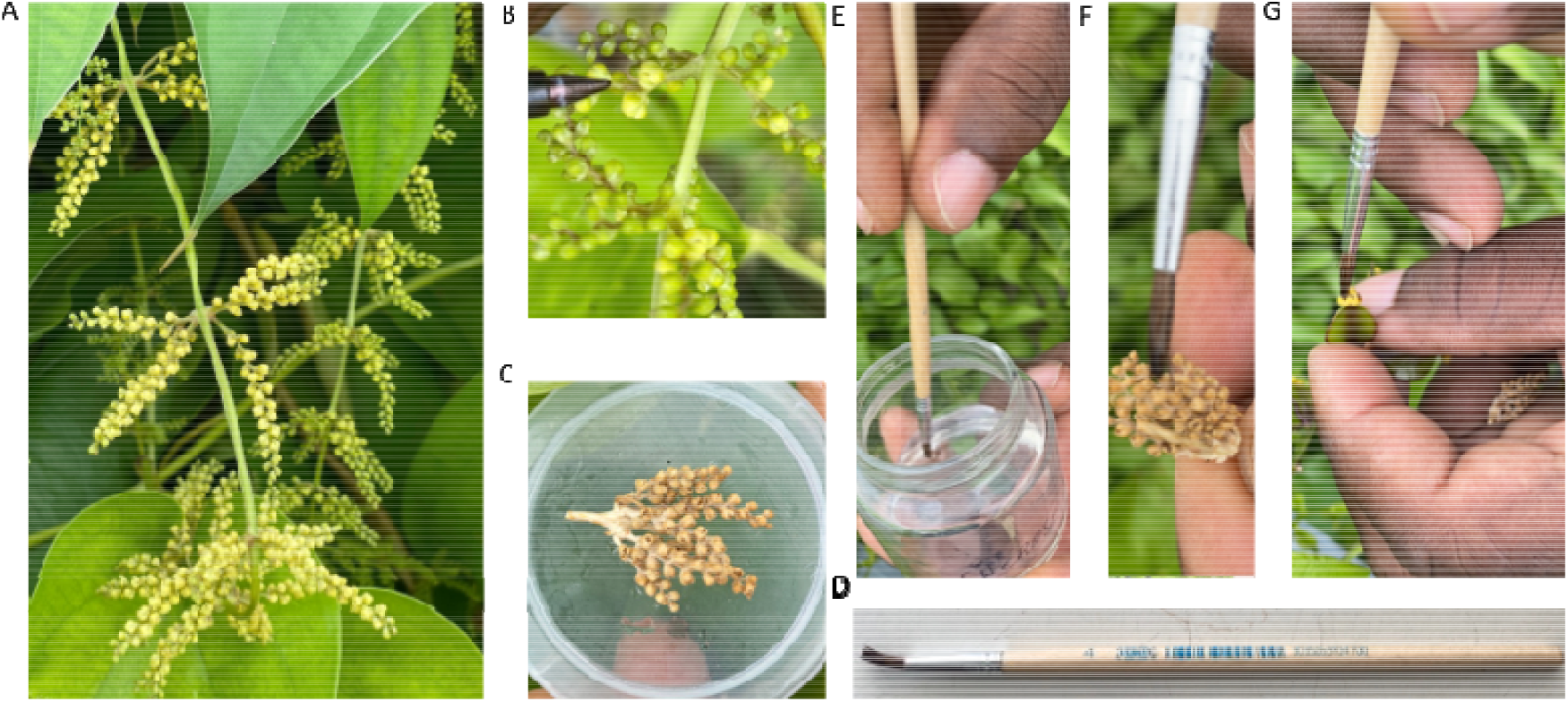
Collection of fresh mature opened male flower samples. A) Photo of the inflorescences of 14M with mature flowers; B) Close-up photo of opened male flowers (14M); C) Photo of lyophilized flower samples (14M); D) Picture of the flat paintbrush used for artificial pollination; E) Wetting the paintbrush in a sucrose solution; (F) Brushing the lyophilized flower spike to collect pollen grains; (G) Pollination of the female flower (74F).

### Fertility and germination test of fresh pollens

The fertility rate of the pollen of the flowers collected from each genotype was checked by a colorimetric test [37]. The anthers were removed from the inflorescence and placed on the slide in a drop of alcohol to remove grease and resin and wiped with Joseph paper (Fisherbrand, grade quality 605). Later, the slides were dried at 35 °C to fix the pollens. After four alcohol-drying wash cycles, Alexander’s staining solution was added. The pollen staining was observed under a microscope (Ernst Leitz Wetzlar GmbH, Germany) with a 400 magnification. Three slides were prepared for each genotype and observed for colored pollens.

In addition, an *in vitro* germination test of the pollen on a synthetic medium was carried out. It consisted in distributing a few drops of the sugar solution containing pollen grains on a Brew-baker and Kwack (BKM) medium composed of 8% of sucrose, 50 mg / L of boric acid (H_3_BO_3_), 300 mg / L of Ca (N0_3_) 2.4H_2_O, 200 mg / L of MgSO_4_.7H_2_O, 100 mg / L of KNO_3_ and 0.7% agar at pH 6.5 [38]. The culture medium was sterilized in an autoclave and poured into three small Petri dishes per genotype.

A minimum of 4 to 5 h of incubation at 25 °C on the medium is necessary in order to observe the germination of the pollens. From 24 h, we determined the percentage of germinated grains by observing with a binocular magnifying glass. A pollen grain was considered germinated *in vitro* if its pollen tube is at least the diameter of the pollen itself [39].

### Pollen lyophilization and conservation

Pollen lyophilization was performed using two sets of treatments. Treatment 1 (T1) consisted of 48 h drying, while treatment 2 (T2) consisted of 72 h drying. The freeze dryer Martin Christ, model Alpha 1-4 (Martin Christ Gefriertrocknungsanlagen GmbH, Germany), was used for the lyophilization. Eppendorf tubes containing flower samples were placed in the freeze dryer. The freezing time was kept for 5 h at −35 °C. After freeze-drying, the samples were placed in a closed bag and stored in a dark jar with silica gel at room temperature 24-25 °C. Freeze-dried pollens were stored for 12 months (Figure 1C).

### *In vitro* fertility and germination test of lyophilized pollens

The lyophilized flower samples were crushed slightly on a slide and placed in a tube containing a sugar solution. After centrifugation, pollen grains were taken and stained with Alexander solution and observed under a microscope, as explained above. Similarly, a germination test, as explained above was carried out. Three replicates per genotype were used.

### Crossing for obtaining hybrid plants from lyophilized pollens

Sixteen tuber fragments of each female genotype were planted in an isolated plot [3]. During the flowering period, some immature inflorescences were covered with paper wrappers. One month after the start of flowering, the female flowers became receptive, and lyophilized pollens from the males were used to pollinate female flowers (74F x 14M and CT198 x Toufi-Tetea) in 2021 and (74F x 14M, CT198 x Toufi-Tetea, 14M x Boutou and CT198 x Ti-violet) in 2022.

In order to facilitate the cross with the lyophilized flower samples, we set up a new technique. A flat paintbrush was immersed in sucrose solution, then used to wet the female flower (Figure 1D). A single male flower was detached from the lyophilized flower spikes and placed on top of a female flower (named: single-male-flower approach). We also tried another technique (named: brushing approach) to collect pollen samples by brushing the lyophilized flower spikes using a wet paintbrush (with a sucrose solution) (Figure 1E-F). A colorimetric test [37] showed that the paintbrush was covered by pollen grains (Data not shown). Then, the wet paintbrush covered by pollen grains can be used to brush and pollinate the female flowers (Figure 1G). Both approaches resulted in successful pollinations.

A total of 100 flowers of each female genotype were crossed using the single-male-flower approach. After crossing, the flowers were protected by the paper wrapper for 2 weeks. We also used fresh pollen samples for pollination. The fruit and seed setting rates were noted two months later. The seeds obtained were germinated on a nutrient medium: M20 (Murashige and Skoog medium with vitamins 3 g, Myoinositol 5 g, sucrose 20 g, Gelrite 2.5 g, activated carbon 1 g, for 1000 ml of the medium; the pH was adjusted to 5.8). The seedlings obtained were propagated *in vitro* and were later acclimatized and planted in pots.

### Genotyping for paternity testing

Leaf samples from five hybrids and their parents (74F x 14M) were used for DNA extraction with the Mixed Alkyl Trimethyl Ammonium Bromide method [40]. DNA quality and quantity were checked on agarose gel and Invitrogen Qubit Flex Fluorometer. A total of 19 microsatellite markers (Table S1) were initially tested for polymorphism between the parents. Seven detected polymorphic markers were used for genotyping the five hybrids. M13 tail (CACGACGTTGTAAAACGAC) was added to the primers. The PCR reaction was performed using the QIAGEN kit under the conditions as followed 95 °C for 5 min, 10 cycles of 30 s at 95 °C, 60 °C for 1 min 30 s and 72 °C for 30 s, 25 cycles of 30 s at 95 °C, Tm °C for 1 min 30 s and 72 °C for 30 s, followed by 30 min at 60 °C. Migration of the PCR products was conducted on the ABI 3500xL (Thermo Fisher Scientific Inc.). Analysis of microsatellite profiles was performed with Genemapper v6.0 (Applied Biosystems™).

### Statistical analysis

GraphPad Prism v9.0.0121 (GraphPad 159 Software Inc., La Jolla, CA, USA) was used for graph construction and statistical analysis. Statistical differences were performed by independent *t*-test at *P* < 0.05.

## Results

### Assessment of fresh pollen viability

A systematic approach was followed to evaluate the effect of lyophilization of pollen grains from the two genotypes viz., 14M and Toufi-Tetea on their viability and suitability for crossbreeding (Figure 2). It is important to ascertain the fertility of pollen grains from the genotypes used for the assay. Hence, fresh pollens were collected from both genotypes and subjected to fertility and germination tests during the first step. The results indicated similar and high pollen fertility (78 and 85% in Toufi-Tetea and 14M, respectively) and pollen germination percent (82 and 86% in 14M and Toufi-Tetea, respectively) in the two genotypes (Figure 3). Therefore, the selected genotypes are good candidates for pollen lyophilization and long-term storage.

**Figure 2.**
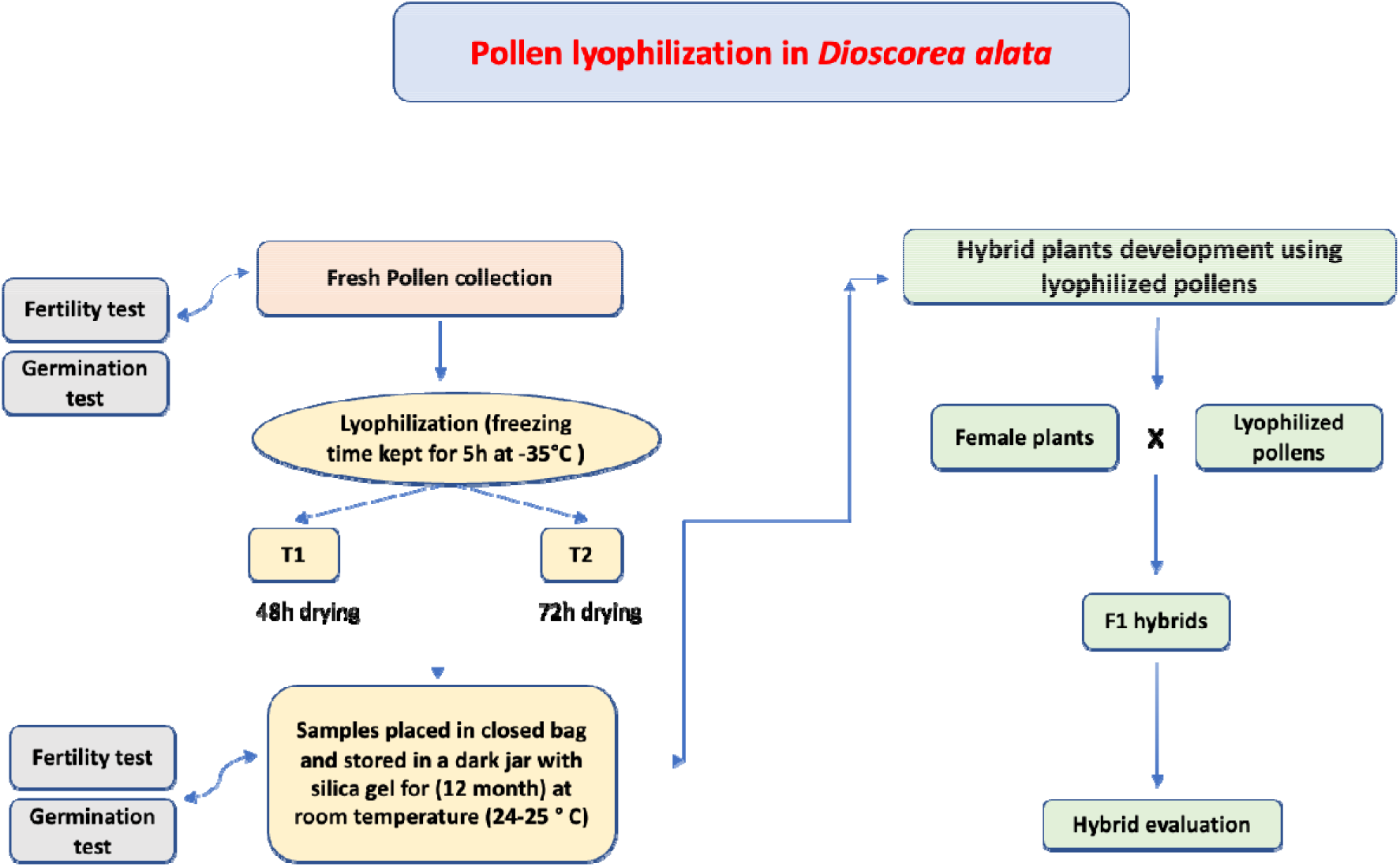
Schematic diagram of hybrid development using lyophilized pollens in *Dioscorea alata.*

**Figure 3.**
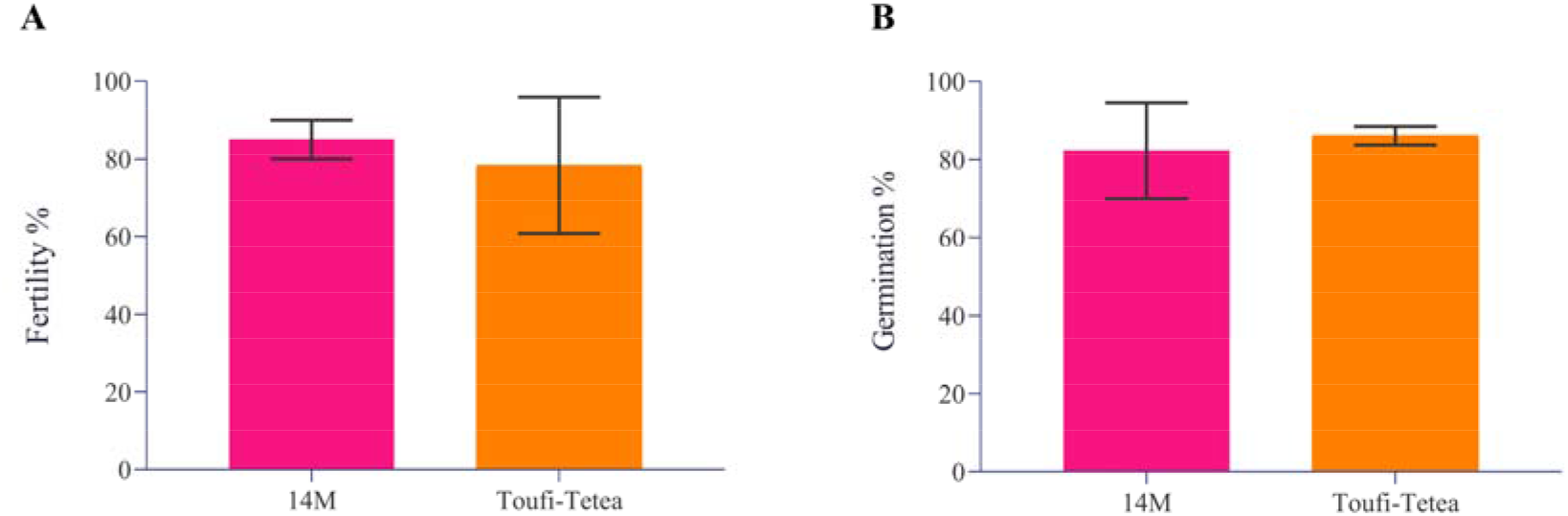
Pollen viability test of two *Dioscorea alata* male varieties used in the current study. A) Fertility percent of freshly collected pollens from two *D. alata* varieties viz., 14M and Toufi-Tetea; B) Germination percentage of freshly collected pollens from two *D. alata* varieties viz., 14M and Toufi-Tetea.

### Lyophilization, storage and their effects on pollen viability

Pollen lyophilization was performed using two treatments (T1 and T2). Both varieties showed a higher germination percent in T1 compared to T2 (Figure 4). The germination percentage was estimated monthly (from November to July) after the pollen lyophilization, and a continuous decrease in germination percentage was observed. 14M showed a decrease in germination percent from 76 to 25% in T1, while a decrease from 50 to 10% under T2 was observed. Similarly, the germination percentage of lyophilized pollens of Toufi-Tetea decreased from 76 to 25% under T1, while under T2, it decreased from 75 to 20%.

**Figure 4.**
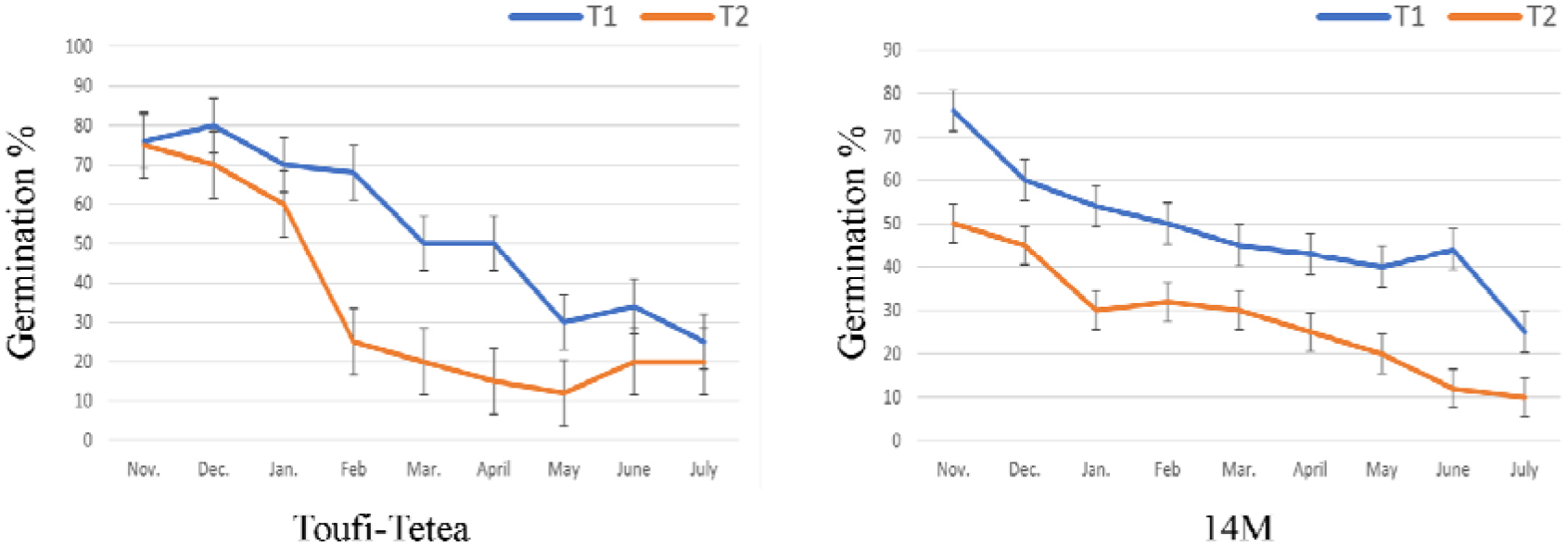
*In-vitro* germination percentage of pollens after lyophilization in A) 14 M and B) Toufi-Tetea from November to July. T1 (48 h drying) and T2 (72 h drying) represents the two lyophilization treatments. Data represent the average of two-year experiments.

The remaining samples of the lyophilized pollens were further stored for 4 more months (12 months in total). A fertility test was performed to check the pollen viability after 12 months of storage. Interestingly, pollens from both varieties displayed a higher fertility percentage in T1 compared to T2 (Figure 5). Altogether, our results showed that T1 (48 h drying) is more conducive for the long-term fertility of pollen grains in *D. alata.*

**Figure 5.**
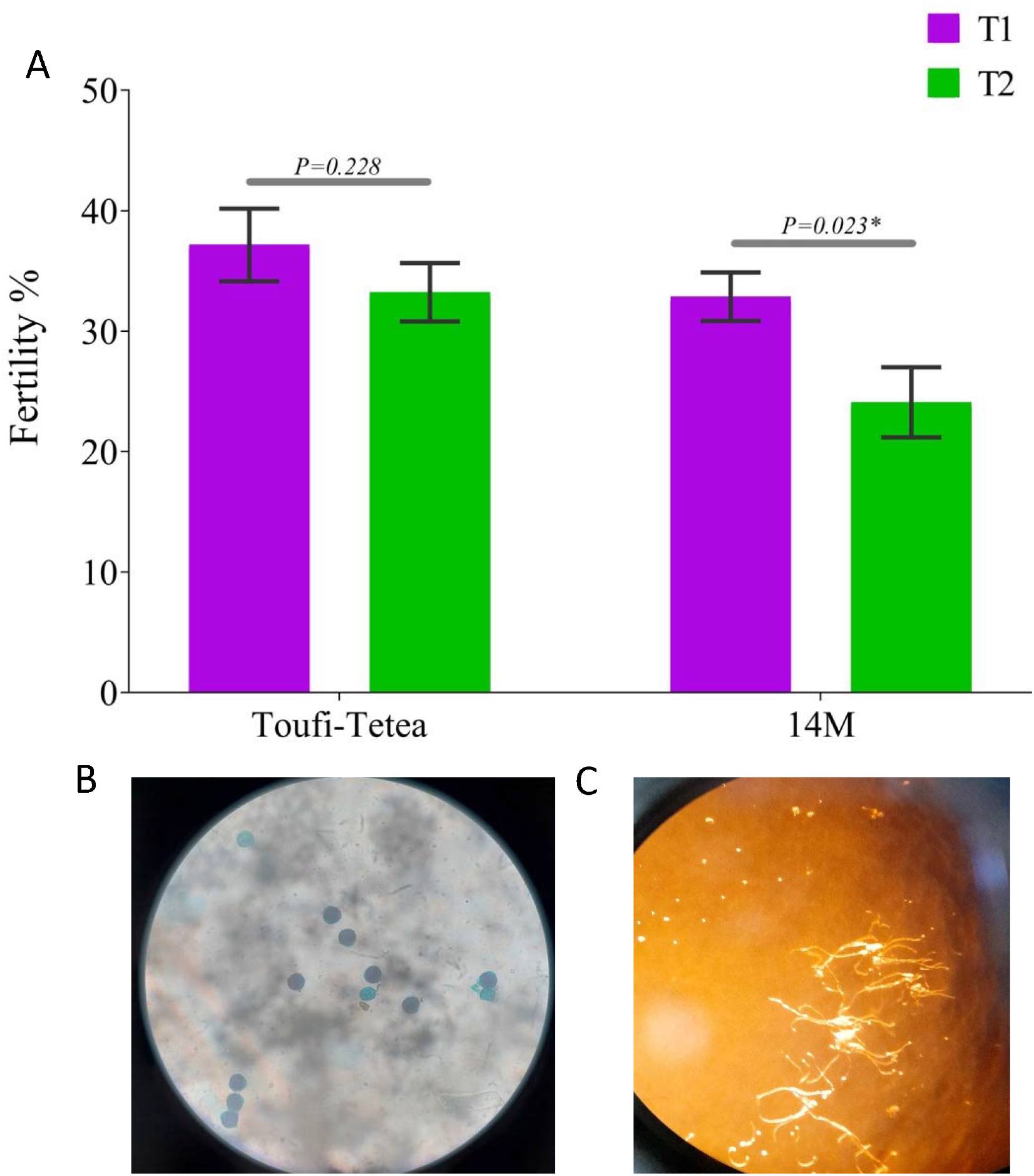
Fertility test of lyophilized pollens after 12 months storage sourced from 14M and Toufi-Tetea. A) Fertility percentage of lyophilized pollens in 14M and Toufi-Tetea; B) Photo of colored lyophilized pollen grains from 14M with the Alexander solution; C) *In vitro* germination of lyophilized pollen grains from 14M. *P* < 0.05 was considered significant (*) using a two-tailed *t*-test.

### Development of hybrid yam plants from lyophilized pollens

To test whether the lyophilized pollens could be used for developing hybrid yam plants after 12 months storage, we crossed the female plants in the field conditions using pollens from T1. Four varieties, 74F, Boutou, Ti-violet and CT-198, were used as female parents, while lyophilized pollens from 14M and Toufi-Tetea were used for pollination. The cross 74F x 14M, CT-198 x Toufi-Tetea, Boutou x 14M, and Ti-violet x Toufi-Tetea yielded 40, 50, 58, and 56% fruits, respectively (Table 1, Figure 6). We also performed crossing using fresh pollens, and the fruit setting rate was higher than lyophilized pollens. The fruits were later evaluated after maturity for viable seeds. Globally, most fruits obtained from lyophilized pollens contain a lot of viable seeds similar to observations with fresh pollens (Table 1).

**Figure 6.**
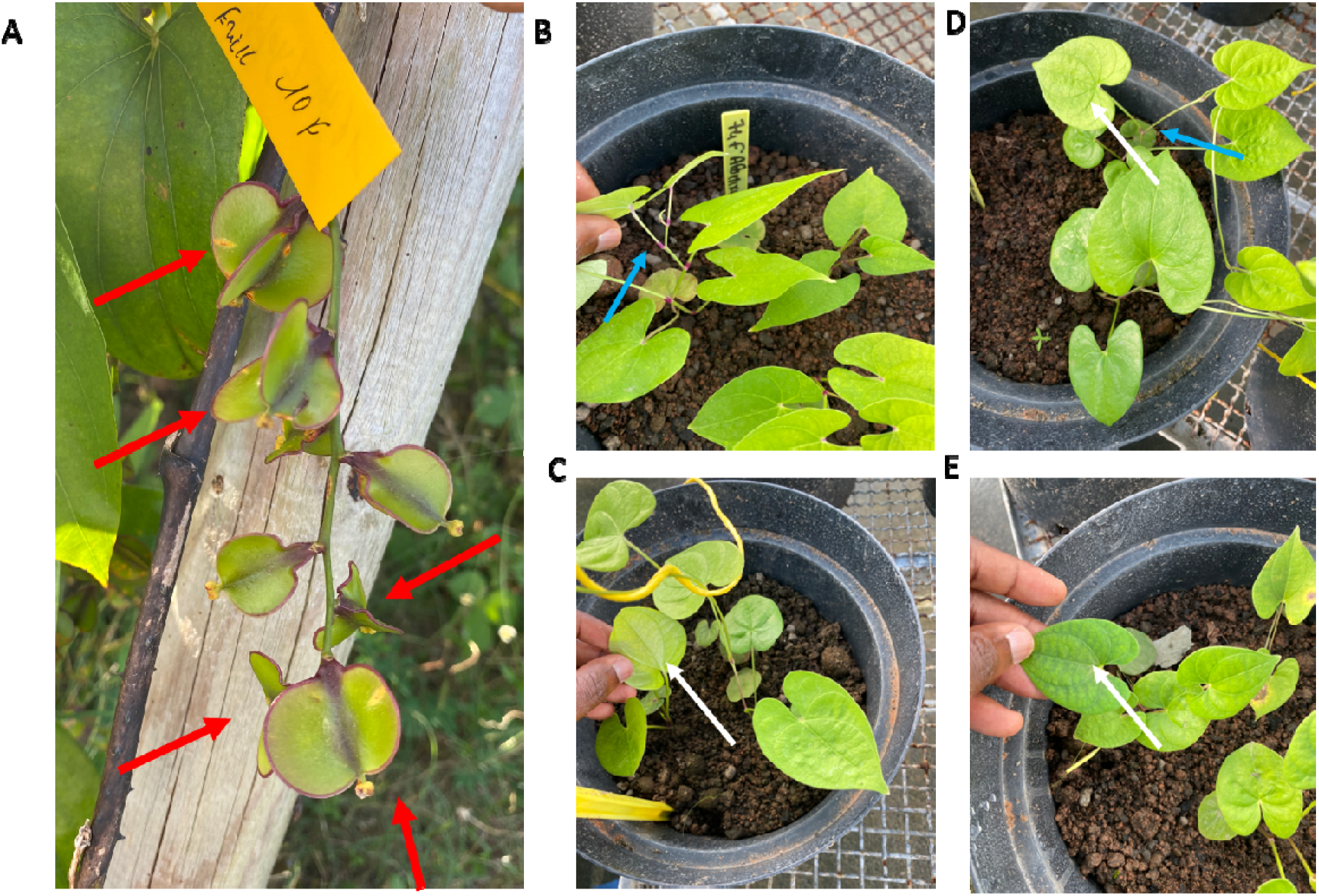
Development of hybrid plants obtained from crosses with lyophilized and 12-month-stored pollens in *Dioscorea alata.* A) Visible swelling ovary of the female flowers 10 days after pollination in 74F x 14M; Photos of seedlings generated from tubers of parents B) 74F, C) 14M, and seeds of (D-E) two selected hybrids obtained from lyophilized pollens and grown in a greenhouse. Red arrows show swollen ovary of the female flowers; blue arrows show the pink coloration present on the petiole of 74F, absent on 14M and segregating in the hybrids. The white arrows show the oval leaf shape of 14M present in both hybrids.

**Table 1.** Development of hybrid plants from lyophilized pollens in *Dioscorea alata*

| Category | Crosses | Number of Flowers pollinated | Number of fruits | Seeds with embryos | Number of plants obtained |
| --- | --- | --- | --- | --- | --- |
| **Lyophilized pollens** | 74F x 14M | 100 | 40 | 20 | 12 * |
|  | CT-198 x Toufi-Tetea | 100 | 50 | 30 | 17 * |
|  | Boutou x 14M | 100 | 58 | 32 | 19 |
|  | Ti-violet x Toufi-Tetea | 100 | 56 | 35 | 22 |
| **Fresh pollens** | 74F x 14M | 100 | 41 | 36 | 21 * |
|  | CT-198 x Toufi-Tetea | 100 | 53 | 44 | 27 * |
|  | Boutou x 14M | 100 | 57 | 30 | 25 |
|  | Ti-violet x Toufi-Tetea | 100 | 66 | 48 | 31 |

\*Data represent the average of two-year experiments.

The obtained viable seeds from the lyophilized pollens were germinated on a nutrient medium (M20), and later seedlings were transplanted in pots. Interestingly, we successfully generated 12-22 hybrid seedlings from the different parent pairs and the seedlings grew normally in a greenhouse (Figure 6).

To further confirm the status of the obtained plants, we performed a paternity testing using seven polymorphic microsatellite markers (between the two parents) on five selected hybrids from the cross 74F x 14M. Several missing data were noted due to the quality of the molecular markers. The genotyping showed a maximum of two alleles for all markers which matches well the ploidy level of the genotypes (Table 2). Overall, the hybrids displayed different allelic combinations from both parents at the different markers (Table 2), confirming that the plants obtained are well from gametes of the two parents.

**Table 2.** Microsatellite markers based paternity testing of five *Dioscorea alata* hybrids (Hyb_1-Hyb_5) obtained from lyophilized and 12-month-stored pollens. Seven polymorphic microsatellite markers between the two parents (14M and 74F) were used (mDaCIRxx) and the alleles (size) were coded as A1/A2. Missing or undetected alleles are coded by “-“. Different colors at each marker represent the different alleles.

| Samples | Names | mDaCIR<br>11_A1 | mDaCIR<br>11_A2 | mDaCIR<br>2_A1 | mDaCIR<br>2_A2 | mDaCIR<br>20_A1 | mDaCIR<br>20_A2 | mDaCIR<br>56_A1 | mDaCIR<br>56_A2 | mDaCIR<br>59_A1 | mDaCIR<br>59_A2 | mDaCIR<br>61_A1 | mDaCIR<br>61_A2 | mDaCIR<br>79_A1 | mDaCIR<br>79_A2 |
| --- | --- | --- | --- | --- | --- | --- | --- | --- | --- | --- | --- | --- | --- | --- | --- |
| Male<br>parent | 14M | 189 | - | 271 | 279 | 202 | 208 | 267 | 271 | 217 | - | 205 | 235 | 255 | 259 |
| Female<br>parent | 74F | 191 | 200 | 259 | 279 | 202 | 208 | 257 | 267 | 220 | - | 217 | - | 254 | 255 |
| Hybrid | Hyb_1 | 200 | - | 259 | 271 | 202 | - | 267 | 271 | 220 | - | 217 | - | 255 | - |
| Hybrid | Hyb_2 | 191 | - | 259 | 279 | 202 | 208 | 257 | 267 | 220 | - | 217 | - | 255 | - |
| Hybrid | Hyb_3 | 189 | 191 | 259 | 271 | 202 | 208 | 257 | 271 | 217 | 220 | 217 | - | 254 | 259 |
| Hybrid | Hyb_4 | 189 | 200 | 259 | 279 | 202 | 208 | 257 | 267 | 217 | 220 | 217 | 235 | 254 | 255 |
| Hybrid | Hyb_5 | 189 | 191 | 259 | 279 | 202 | - | 267 | - | 220 | - | 217 | 235 | 255 | - |

In brief, our study revealed that long-term conserved lyophilized pollens could be used to develop hybrid plants in *D. alata*.

## Discussion

Yam is a significant food crop in developing countries feeding over 150 million people [41]. A fast genetic improvement of yams is required in order to satisfy the growing demand of yam tubers owing to the rise of demography coupled with the negative effects of climate change on agriculture. Low tuber yield and nutritional quality as well as the plant susceptibility to biotic (anthracnose and viruses) and abiotic stresses are the major impediments to yam production worldwide [14]. Unfortunately, various biological constraints limit yam improvement [12–16]. It is necessary to overcome these problems with limited resources to achieve successful breeding programs and accelerate genetic gains. Hereby, we report a practicable protocol for the long-term storage of pollen grains and the first successful development of hybrids of *D. alata* from lyophilized pollens.

Lyophilization is a valuable technique for long-term pollen preservation and has the key advantage that the lyophilized samples can be stored at room temperature without any sophisticated equipment. Pollen lyophilization has been used for many crops with genetic and biological constraints, such as Eucalyptus [32], maize [33,34], and pigeon pea [35]. However, the success rates vary according to the plant species. Many studies have reported *in vitro* germination of preserved pollens of yam using different techniques [20,23,24,42–44]. For instance, Mondo et al., [20] achieved 26.4–59.7% *in vitro* germination in *D. alata.* Our study observed a 45% decrease in pollen germination from fresh pollens to one-year lyophilized pollens. Daniel et al., [44] also observed a sharp decrease in pollen germination of yam species after cryo-preservation. They recommended that wet-cold storage (hermetic cold storage without previous drying) is a superior method for long-term pollen viability compared to air-dried storage and freeze-drying. However, none of these studies used long term stored pollens for pollination and hybrid development. As previously demonstrated, the decrease in pollen germination can be attributed to the storage conditions and storage time [45,46]. Mathad et al., [47] observed variable germination rates, fruit set, and seed development of cryo-preserved hot pepper pollens under different temperatures. Moreover, our results depicted significant differences between the two treatments used for lyophilization. Treatment 1 with 48 h drying appeared superior to Treatment 2 with 72 h drying. The low efficacy of T2 may be attributed to the reduced moisture content due to longer drying period [44].

A key objective of this study was achieved with the development of hybrids using long-term lyophilized pollens. Fruit and seed setting rates in tetraploid and diploid varieties using lyophilized pollens were almost similar but we observed that they were lower than with fresh pollens. Likewise, a previous study reported higher fruit and seed setting rates using fresh pollens under field conditions [48]. It is well known that a successful fruit and seed setting is highly dependent on the genotype, environment and requires a precise timing [49]. Nonetheless, our results signify that the normal fruit setting and seed development are achievable using lyophilized pollens in yam breeding. Furthermore, no morphological abnormalities were observed in hybrids developed from lyophilized pollens, in accordance with observations in Eucalyptus [32], maize [33,34], and pigeon pea [35].

Seed setting occurred to a certain extent despite the decrease in pollen viability over time in our study, suggesting that lyophilization is a potential approach for long-term pollen storage to develop hybrids in yam breeding programs. With the availability of lyophilized pollen samples when female flowers become receptive, breeders can overcome the asynchronous flowering issue between male and female *D. alata* genotypes and make cross on a large quantity of female flowers. We would recommend a large quantity of lyophilized and stored pollens to increase the chances of obtaining enough offspring. Nonetheless, the number of hybrid plants generated from lyophilized pollens in this study was quite similar to those from fresh pollens.

The technique developed in this study will facilitate exchanges of pollen samples and increase the available gene pools among breeding programs in different countries and regions of the world. Pollen samples could be sent and stored at ambient temperature which is very useful for labs in developing countries. In future, more genotypes will be tested for pollen storage using the lyophilization approach with an improved protocol to achieve the maximum viability of stored pollens. Although this study was focused on *D. alata,* we expect that our protocol is applicable to other Dioscorea species since a high similarity of floral biology among yam species has been reported [21,49].

## Supporting information

Primer sequences and details of the microsatellite markers used in this study

## Declarations

### Ethics approval and consent to participate

Not applicable

### Consent for publication

Not applicable

### Availability of data and material

All data generated or analysed during this study are included in this published article and its supplementary information file.

### Authors’ contributions

*Conceptualization:* Erick Malédon, Komivi Dossa, Hâna Chair. *Experiment:* Erick Malédon, Komivi Dossa, Elie Nudol, Christophe Perrot, Marie-claire Gravillon, Ronan Rivaillan. *Data acquisition, curation and formal analysis:* Erick Malédon, Komivi Dossa, Elie Nudol, Christophe Perrot, Marie-claire Gravillon, Ronan Rivallan, Denis Cornet, Hâna Chair. *Funding acquisition:* Denis Cornet, Komivi Dossa. *Writing ± original draft:* Erick Malédon, Komivi Dossa. *Writing ± review & editing:* Denis Cornet, Hâna Chair. All authors have read and approved the final version of this manuscript.

## Acknowledgements

Not applicable

## Supplementary files

**Table S1**. Primer sequences and details of the microsatellite markers used in this study.

